# Cold-induced muscle atrophy in zebrafish: Insights from swimming activity and gene expression analysis

**DOI:** 10.1101/2023.09.03.553941

**Authors:** Daisuke Ikeda, Seina Fujita, Kaito Toda, Yuma Yaginuma, Nobuhiro Kan-no, Shugo Watabe

## Abstract

The investigation into the effects of cold acclimation on fish skeletal muscle function and its potential implications for muscle atrophy is of great interest to us. This study examines how rearing zebrafish at low temperatures affects their locomotor activity and the expression of genes associated with muscle atrophy. Zebrafish were exposed to temperatures ranging from 10 °C to 25 °C, and their swimming distance was measured. The expression levels of important muscle atrophy genes, Atrogin-1 and MuRF1, were also evaluated. Our findings show that swimming activity significantly decreases when the water temperature ranges from 10 °C to 15 °C, indicating a decrease in voluntary movement. Additionally, gene expression analysis shows a significant increase in the expression of Atrogin-1 and MuRF1 at 10 °C. This up-regulation could lead to muscle atrophy caused by decreased activity in cold temperatures. To investigate the effects of exercise on reducing muscle atrophy, we subjected zebrafish to forced swimming at a temperature of 8 °C for ten days. This treatment significantly reduced the expression of Atrogin-1 and MuRF1, emphasizing the importance of muscle stimulation in preventing muscle atrophy in zebrafish. These findings suggest that zebrafish can serve as a valuable model organism for studying muscle atrophy and can be utilized in drug screening for muscle atrophy-related disorders. Cold-reared zebrafish provide a practical and ethical approach to inducing disuse muscle atrophy, providing valuable insights into potential therapeutic strategies for addressing skeletal muscle atrophy.

## 1. Introduction

It is widely recognized that fish, being ectothermic animals, exhibit reduced mobility at low water temperatures. To elucidate the mechanism of cold acclimation in medaka (*Oryzias latipes*) skeletal muscle, we performed RNA-Seq analysis and identified temperature-dependent changes in gene expression [1]. Among these genes, the expression level of Atrogin-1 increased by approximately 30-fold in the experimental group raised at a low temperature. Two ubiquitin E3 ligases specific to muscle, Atrogin-1 and MuRF1, have been extensively studied and they play a critical role in skeletal muscle atrophy [2–4]. These genes have been found to be upregulated in disuse muscle atrophy in humans resulting from bed rest or immobilization (such as a cast or leg brace), as well as in the atrophied skeletal muscles of astronauts (reviewed in [4]). Gracey et al. discovered a significant increase in the expression levels of Atrogin-1 in muscle tissues of carp that were acclimatized to 10 °C as compared to the ones acclimatized to 30 °C [5]. They proposed that decreased fish movement at cold temperatures might result in muscle atrophy. We hypothesized that medaka reared in a cold environment and spending most of their time on the bottom of the aquarium without any active movement would experience muscle atrophy [1]. However, the observed phenomenon of “inactivity induced by cold rearing” lacks quantifiable outcomes and remains subjective. Zebrafish (*Danio rerio*) has been extensively studied as a model organism, and its complete genome sequence is publicly accessible [6]. Experiments examining the influence of rearing temperature on zebrafish activity have demonstrated a decrease in maximum sustained swimming speed, number of actions, and activity levels at lower temperatures (reviewed in [7]). However, few studies have quantified the distance swum per unit of time. More recently, Morgan et al. acclimated zebrafish to 15 temperatures ranging from 10 to 38 °C for one month and analyzed changes in several phenotypes, including behavior [8]. They observed that the migration distance (activity, measured in body lengths per minute) decreased at lower temperatures compared to the optimal water temperatures, which are typically around 28 °C.

In this study, we measured the swimming distance of zebrafish at different rearing temperatures and investigated the relationship between their locomotor activity and the expression levels of genes associated with muscle atrophy. Additionally, we investigated whether exercise could suppress muscle atrophy in zebrafish that had reduced voluntary movements as a result of cold rearing.

## Materials and methods

### 2.1. Zebrafish care

Adult zebrafish specimens were obtained from a vendor (Meito-Suien, Aichi, Japan) at an approximately equal sex ratio and were maintained at 28 °C with a 14:10-hour light-dark photoperiod. The fish were hand-fed daily with Hikari Lab 450 micro pellet food (Kyorin, Hyogo, Japan) until they were full. All animal experiments were conducted in compliance with the regulations set by the Animal Experimentation Committee of the School of Marine Biosciences, Kitasato University.

### 2.2. The effects of low temperature

A total of 60 individuals (weight 0.38 ± 0.09 g) were placed in a 60 L commercial tank at 25 °C and reared for 1 week. The water temperature for rearing the zebrafish was gradually decreased in 5 °C increments until it reached 20 °C, 15 °C, and 10 °C before being increased back to 25 °C under the same conditions. The water temperature adjustment took place gradually over three days, with a controlled rate of change limited to 1-2 °C per day. The behavior of a single fish was recorded using a time-lapse camera (TLC200Pro, Brinno Inc., Taipei, Taiwan) at 1-second intervals in a 40 cm x 25 cm tank with a water level of 10 cm at each temperature (n=12). The camera was positioned directly overhead the tank. The video recording was conducted over three consecutive days at each water temperature, with four individuals recorded daily. The recording sessions were intentionally scheduled between 10:00 and 15:00 to align with the circadian rhythms of zebrafish locomotor activity. To investigate the effects of low temperature on zebrafish behavior, the rearing period at 10 °C was extended to 18 days, and the behavior of four individuals was recorded daily. The fish were allowed to adjust to the tank for 10-15 minutes before beginning the 30-minute video recording. The fish were returned to their original tank after being recorded on video. We utilized Kinovea, a free 2D motion analysis software, to measure the distance moved by each individual during a 30-minute period (Video S1-4). The results were visualized using a boxplot created with the ggplot2 package in R. Total RNA was extracted from the dorsal body muscles of the four test fish after conducting a swimming activity test at each water temperature. Following the achievement of the designated temperature of 10 °C, samples were collected from four individuals at predetermined time intervals, specifically on days 18, 21, 24, 27, 30, and 33. The schedule for rearing water temperature changes, video recording, and skeletal muscle sampling is shown in Fig. S1.

### 2.3. The effect of forced swimming

A total of 12 individuals (weight 0.42 ± 0.10 g) were placed in a 60 L tank and reared at 28 °C for 1 week. The tank was divided into compartments with holes to allow water flow, and each compartment housed six fish. The water temperature was then gradually lowered to 8 °C over a 10-day period, decreasing by 2 °C per day. The temperature was then maintained for an additional 2 weeks. The fish in one compartment were moved to a circular tank (Fig. S2, Video S5, S6) with a flow rate set to the maximum speed at which the fish could swim continuously. They were allowed to exercise for 5 hours per day at a temperature of 8 °C for a period of 10 days. The fish in the other compartment did not receive any exercise. Additionally, six control fish reared at 28 °C (weight 0.38 ± 0.08 g) were used. Total RNA was extracted from the dorsal body muscles of all test fish.

### 2.4. RNA extraction and real-time PCR analysis

Total RNA was extracted using ISOGEN II (Nippon Gene, Tokyo, Japan). The extracted RNA was reverse transcribed into cDNA using the ReverTra Ace qPCR RT Master Mix (TOYOBO, Osaka, Japan). The cDNA was analyzed using real-time PCR with THUNDERBIRD SYBR qPCR Mix (TOYOBO) and a LightCycler 96 instrument (Roche, Basel, Switzerland), following the manufacturer’s instructions. The relative expression levels of genes responsible for muscle atrophy were determined using RT-qPCR, with EF1-α serving as the reference gene. The primer sets used in this study are listed in Supplementary Table S1.

### 2.5. Statistical analysis

Student’s t-tests were used to show the statistically significant difference in distance moved at each water temperature compared to the initial temperature of 25 °C at the beginning of the experiment. The effect of four temperatures on the distance moved was analyzed using a one-way ANOVA, followed by a Tukey-Kramer post-hoc test. Additionally, for the qPCR data, a one-way ANOVA followed by a Tukey-Kramer post-hoc test was used. The statistical analysis was performed using R Statistical Software (version 4.2.2). A p-value below 0.05 indicates a statistically significant difference.

## 3. Results and discussion

### 3.1. Effect of low temperature on swimming activity and gene expression related to muscle atrophy

At temperatures of 25 °C and 20 °C, zebrafish moved approximately 40 meters in 30 minutes under both conditions (Fig. 1A, Video S1, S2). However, at 15 °C, their movement decreased significantly to approximately 20 meters in 30 minutes, and this level remained the same at 10 °C (Fig. 1A, Video S3, S4). Even after being reared at 10 °C for 18 days, the movement did not decrease any further (Fig. 1A). When the water temperature was raised to 15 °C, a significant increase in the distance moved was observed. The average distance moved at 20 °C and 25 °C returned to approximately 40 meters in 30 minutes, which was the same level observed at the beginning of the experiment.

**Figure 1.**
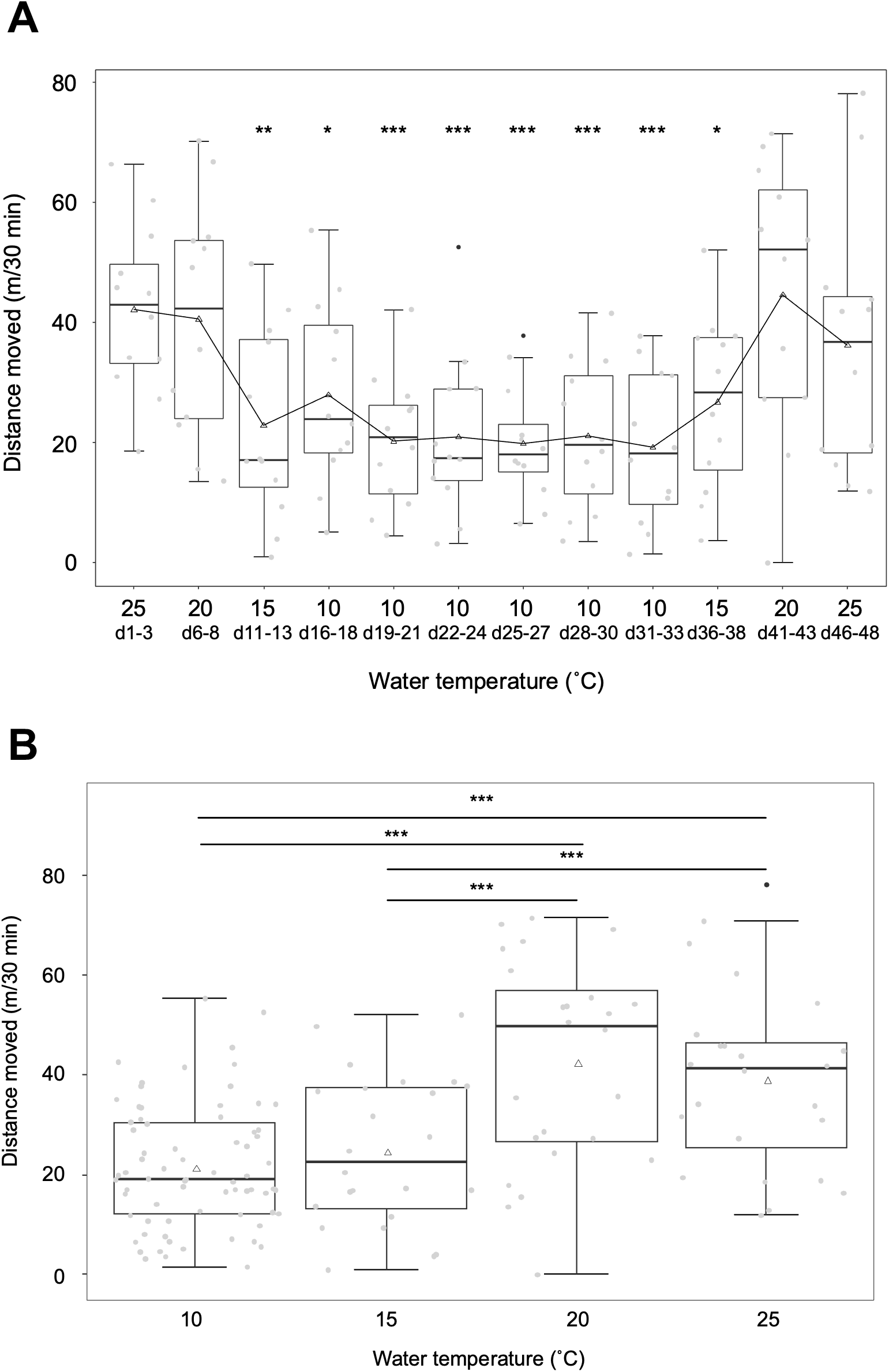
The relationship between temperature and swimming activity of zebrafish. A box plot was used to illustrate the distances moved by zebrafish in each experimental group. Black dots indicate outliers. Triangle markers represent the average distances moved. A. Changes in swimming activity in response to temperature variations. Voluntary swimming activity decreases with lower temperatures but increases as temperatures rise. Asterisks indicate significant differences compared to swimming activities at the initial water temperature of 25 °C ^(*:^ p < 0.05, ^**:^ p < 0.01, ^***:^ p < 0.001). B. Swimming activity at different temperatures. A significant difference in distance moved was observed between temperatures of 20 °C and 15 °C. In the post-hoc test, asterisks are used to indicate statistical significance in comparisons between swimming activities at different temperatures ^(***:^ p < 0.001).

We compared the distance moved at different water temperatures, disregarding the number of acclimation days. Our focus was solely on determining the effect of water temperature on zebrafish swimming activity (Fig. 1B). The results show no significant difference in distance moved at temperatures of 10 and 15 °C, as well as at temperatures of 20 and 25 °C (Fig. 1B). A notable disparity in distance moved was observed between the temperature ranges of 10-15 °C and 20-25 °C (Fig. 1B).

The study by Morgana et al. found that swimming activity, measured in body lengths moved per minute, decreased from 200 to 100 when comparing temperatures of 22 °C and 14 °C, indicating a decrease of approximately 50% [8]. It has been reported that when water temperature drops below 15 °C, there is a significant decrease in swimming activity and noticeable changes in normal behavioral patterns [9]. Wakamatsu et al. found that zebrafish reared at 28 °C experienced a significant decrease in critical swimming speed when the water temperature dropped from 16 °C to 14 °C [10]. The observations indicate that zebrafish activity significantly decreases within the temperature range of 20 °C to 15 °C.

The expression levels of Atrogin-1 and MuRF1, which are the main muscle-specific E3 ubiquitin ligase genes responsible for muscle atrophy, including sarcopenia [2–4], were comparable at both 25 °C and 20 °C. However, at 10 and 15 °C, the expression levels of Atrogin-1 and MuRF1 were higher compared to 25 °C at the start of the experiment, with Atrogin-1 levels being 6 to 10 times higher and MuRF1 levels being 20 to 80 times higher (Fig. 2A). The expression levels of the genes decreased as the water temperature increased, and there were no differences observed from the beginning of the experiment at either 25 °C or 20 °C. No significant changes in gene expression levels were observed for Cbl-b and Nedd4 [3,11], which have been linked to muscle atrophy, or for myosin heavy chain (Myhz2), the primary protein responsible for muscle contraction and known to be upregulated during exercise-induced growth [12] (Fig. S3).

**Figure 2.**
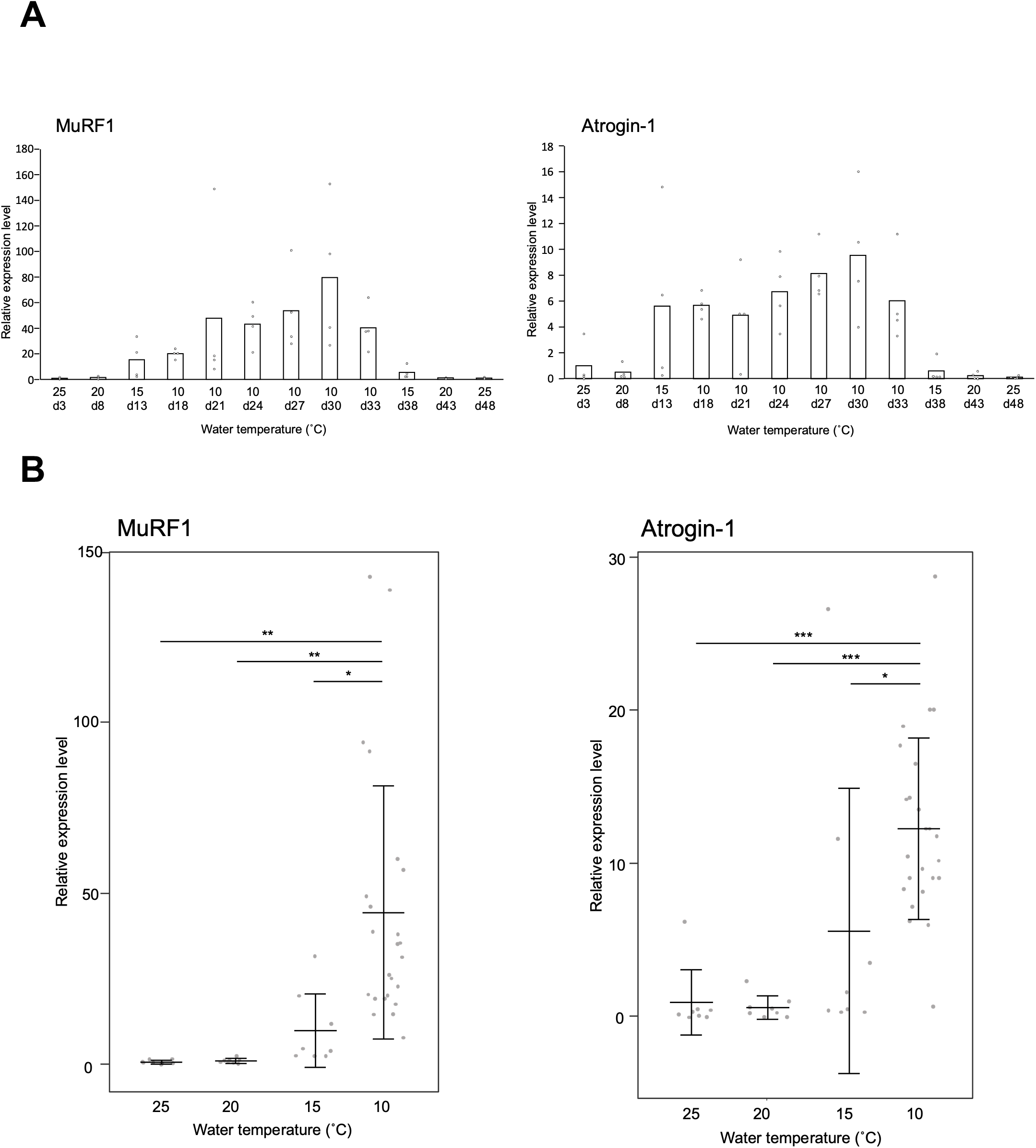
The relationship between temperature and the expression levels of MuRF1 and atrogin-1 in zebrafish muscle tissues. The relative expression levels were normalized using EF-1α as a reference and presented as a ratio to the initial water temperature of 25 °C (A; n=4) and the water temperature of 25 °C (B; n=8). The expression levels are presented as means (A; n=4) and means ± standard deviations (B; 25, 20, and 15 °C: n=8; 10 °C: n=24). In the post-hoc test, asterisks indicate statistical significance in comparisons between expression levels at different temperatures (^*:^ p < 0.05, ^**:^ p < 0.01, ^***:^ p < 0.001).

To specifically examine the impact of water temperature on gene expression levels related to muscle atrophy, we compared the gene expression levels at each water temperature (Fig. 2B). Statistically significant differences in gene expression levels were observed at 10 °C compared to expression levels at 20 °C, 25 °C, and even at 15 °C for both genes (Fig. 2B). The gene expression levels at 15 °C were higher than those at 20 °C and 25 °C, but the differences were not statistically significant. There was no significant difference in the activity level of zebrafish at 10 and 15 °C (Fig. 1B). The expression level of genes associated with muscle atrophy was significantly higher at 10 °C (Fig. 2B), suggesting a higher incidence of muscle atrophy. Additionally, the gene expression levels of Cbl-b and Nedd4 were found to be slightly higher only at 10 °C (Fig. S3). These findings indicate that factors other than decreased physical activity contribute to muscle atrophy during low-temperature rearing. However, further experiments are necessary to accurately assess the extent of muscle atrophy at temperatures of 10 °C and 15 °C. These experiments should include matching rearing periods and sample sizes. Muscle atrophy can occur due to fasting or inadequate nutritional intake [13]. In research conducted on fish, it has been observed that fasting induces upregulation of Atrogin-1 and MuRF1 in zebrafish [14], Atlantic salmon [15,16], and rainbow trout [17]. It would have been beneficial to evaluate food consumption in both temperature zones; however, this aspect was not carried out in the conducted experiment.

### 3.2. Effect of forced swimming on gene expression related to muscle atrophy

Cold rearing reduced the activity level of zebrafish and was associated with an upregulation of genes associated with muscle atrophy, indicating that muscle atrophy occurrs by disuse. We hypothesized that stimulating the muscles of cold-reared zebrafish through forced swimming could suppress the expression of genes associated with muscle atrophy. The expression levels of Atrogin-1 and MuRF1 in zebrafish reared at 28 °C were compared to those reared at 8 °C. Atrogin-1 expression increased by 9.4-fold, while MuRF1 expression increased by 42.7-fold (Fig. 3). Subjecting the zebrafish to 5 hours of daily forced swimming over 10 consecutive days resulted in a significant reduction in gene expression levels. Both genes exhibited a decrease to about 25% of their expression levels in the non-exercised condition (Fig. 3). The data strongly suggest that reduced voluntary movements, primarily caused by cold rearing, are a significant contributor to muscle atrophy in zebrafish. Exercise training is a highly effective way to combat disuse atrophy. Several factors, such as endurance and resistance exercise, exercise duration, and the age and sex of the individual, have been shown in experiments with humans and rodents to affect the prevention and recovery of disuse muscle atrophy (reviewed in [4]). Additionally, exercise training improves the swimming ability of zebrafish during their juvenile stage (8-12 months) and middle-aged stage (15-20 months), but not in their old age (25-30 months) [18]. The zebrafish used in this study are juveniles, approximately 6-8 months. It is important to acknowledge that using older individuals may yield different results. Additionally, studies have shown that excessive exercise can result in muscle atrophy and an increase in the expression of Atrogin-1 and MuRF1 in zebrafish muscle [19]. Cold-reared zebrafish can be valuable for studying the ideal intensity and duration of exercise needed to prevent muscle atrophy.

**Figure 3.**
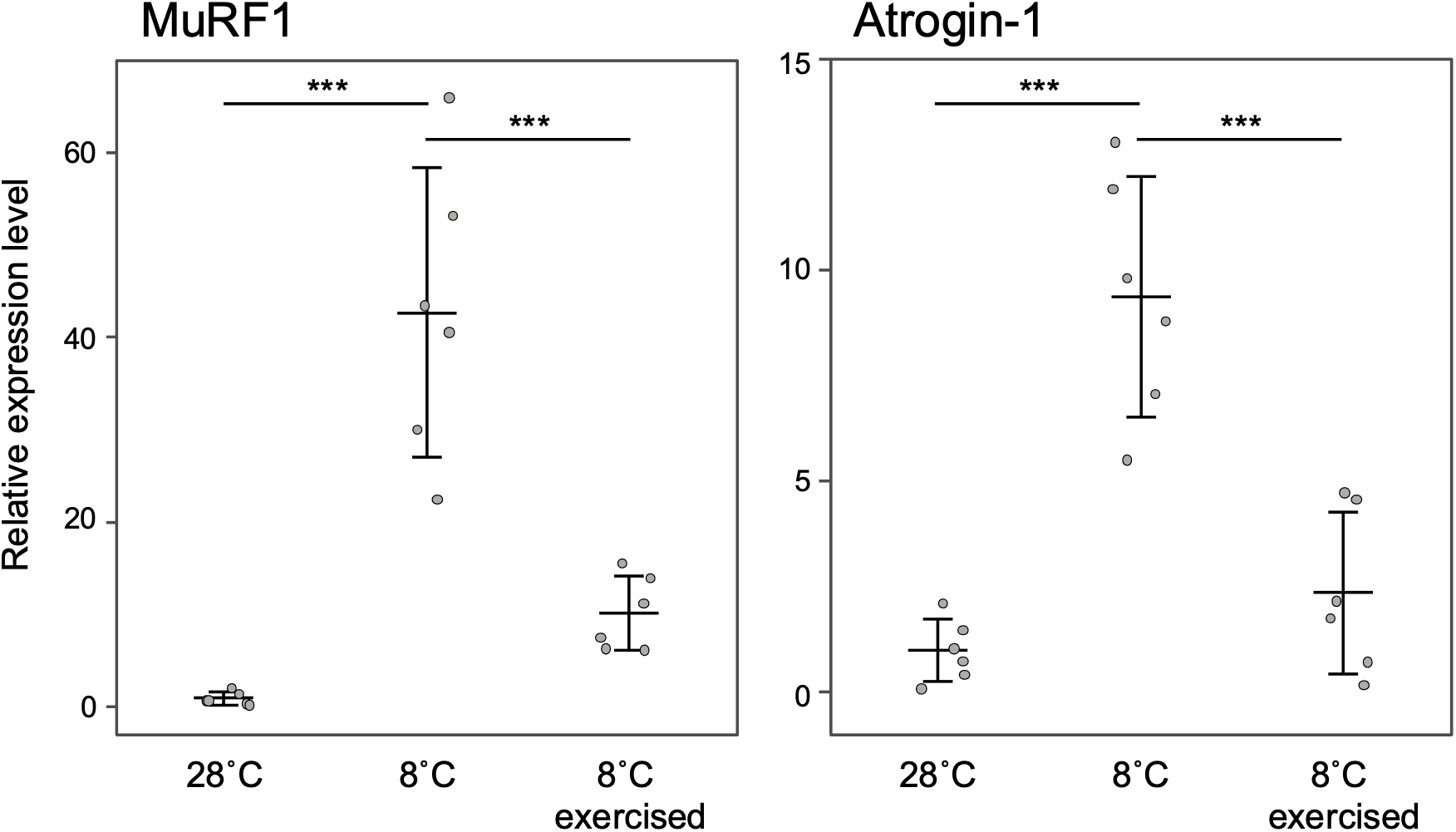
The relationship between forced exercise and the expression levels of MuRF1 and atrogin-1 in low temperature rearing. The relative expression levels were normalized using EF-1α as a reference and presented as a ratio to the levels at a water temperature of 28 °C. The expression levels are presented as means ± standard deviations (n=6). In the post-hoc test, asterisks indicate statistical significance in comparisons between expression levels under different conditions (^***:^ p < 0.001).

In rodent models, muscle atrophy can be induced through artificial methods such as hind limb suspension, nerve transection, and cast immobilization [20,21]. The study findings strongly suggest that zebrafish can experience muscle atrophy due to disuse when kept in low temperatures. This approach seems to offer benefits in terms of efficiency, cost, and ethical considerations. Zebrafish have been proposed as a model for studying sarcopenia due to its ability for rapid genetic intervention and its conserved genetic and physiological features [22]. Additionally, zebrafish have been utilized as model organisms for large-scale drug screening [23], including in the treatment of muscle diseases [24–26]. Utilizing cold-reared zebrafish for drug screening could provide valuable insights into developing therapeutics to prevent muscle atrophy. This study demonstrates the usefulness of zebrafish as a model organism for studying muscle atrophy and emphasizes its potential for drug screening in muscle-related disorders. The use of zebrafish reared in a cold environment offers an ethical and practical method to induce disuse muscle atrophy. This approach provides valuable insights into potential therapeutic strategies for diseases related to muscle atrophy.

More recently, in a study on sarcopenia using zebrafish as a model, Rutkove et al. examined the distance traveled per unit of time by young (6 months) and old (33 months) fish [27]. The results show that older fish travel approximately half the distance of younger fish. Although securing a sufficient number of old fish for sarcopenia studies is challenging, our results suggest that the simple method of low-temperature rearing could allow for the use of young fish as a substitute for older ones. Comparing muscle atrophy in cold-reared and aged zebrafish is an intriguing area for future research.

## References

[1] D. Ikeda, H. Koyama, N. Mizusawa, N. Kan-no, E. Tan, S. Asakawa, S. Watabe, Global gene expression analysis of the muscle tissues of medaka acclimated to low and high environmental temperatures, Comp. Biochem. Physiol. Part D Genomics Proteomics. 24 (2017) 19–28. 10.1016/j.cbd.2017.07.002.

[2] J.P. Gumucio, C.L. Mendias, Atrogin-1, MuRF-1, and sarcopenia, Endocrine. 43 (2013) 12–21. 10.1007/s12020-012-9751-7.

[3] D. Peris-Moreno, L. Cussonneau, L. Combaret, C. Polge, D. Taillandier, Ubiquitin Ligases at the Heart of Skeletal Muscle Atrophy Control, Molecules. 26 (2021) 407. 10.3390/molecules26020407.

[4] Y. Gao, Y. Arfat, H. Wang, N. Goswami, Muscle Atrophy Induced by Mechanical Unloading: Mechanisms and Potential Countermeasures, Front. Physiol. 9 (2018). 10.3389/fphys.2018.00235.

[5] G.B. McClelland, P.M. Craig, K. Dhekney, S. Dipardo, Temperature- and exercise-induced gene expression and metabolic enzyme changes in skeletal muscle of adult zebrafish (Danio rerio), J. Physiol. 577 (2006) 739–751. 10.1113/jphysiol.2006.119032.

[6] K. Howe, M.D. Clark, C.F. Torroja, J. Torrance, C. Berthelot, M. Muffato, J.E. Collins, S. Humphray, K. McLaren, L. Matthews, S. McLaren, I. Sealy, M. Caccamo, C. Churcher, C. Scott, J.C. Barrett, R. Koch, G.-J. Rauch, S. White, W. Chow, D.L. Stemple, The zebrafish reference genome sequence and its relationship to the human genome, Nature. 496 (2013) 498–503. 10.1038/nature12111.

[7] J.F. López-Olmeda, F.J. Sánchez-Vázquez, Thermal biology of zebrafish (Danio rerio), J. Therm. Biol. 36 (2011) 91–104. 10.1016/j.jtherbio.2010.12.005.

[8] R. Morgan, A.H. Andreassen, E.R. Åsheim, M.H. Finnøen, G. Dresler, T. Brembu, A. Loh, J.J. Miest, F. Jutfelt, Reduced physiological plasticity in a fish adapted to stable temperatures, Proc. Natl. Acad. Sci. 119 (2022) e2201919119. 10.1073/pnas.2201919119.

[9] R.L. Malek, H. Sajadi, J. Abraham, M.A. Grundy, G.S. Gerhard, The effects of temperature reduction on gene expression and oxidative stress in skeletal muscle from adult zebrafish, Comp. Biochem. Physiol. Part C Toxicol. Pharmacol. 138 (2004) 363–373. 10.1016/j.cca.2004.08.014.

[10] Y. Wakamatsu, K. Ogino, H. Hirata, Swimming capability of zebrafish is governed by water temperature, caudal fin length and genetic background, Sci. Rep. 9 (2019) 16307. 10.1038/s41598-019-52592-w.

[11] T. Nikawa, K. Ishidoh, Ubiquitin ligase Cbl-b and inhibitory Cblin peptides, Biochim. Biophys. Acta BBA - Proteins Proteomics. 1868 (2020) 140495. 10.1016/j.bbapap.2020.140495.

[12] A.P. Palstra, C. Tudorache, M. Rovira, S.A. Brittijn, E. Burgerhout, G.E.E.J.M. van den Thillart, H.P. Spaink, J.V. Planas, Establishing Zebrafish as a Novel Exercise Model: Swimming Economy, Swimming-Enhanced Growth and Muscle Growth Marker Gene Expression, PLOS ONE. 5 (2010) e14483. 10.1371/journal.pone.0014483.

[13] S. Cohen, J.A. Nathan, A.L. Goldberg, Muscle wasting in disease: molecular mechanisms and promising therapies, Nat. Rev. Drug Discov. 14 (2015) 58–74. 10.1038/nrd4467.

[14] I.P.G. Amaral, I.A. Johnston, Insulin-like growth factor (IGF) signalling and genome-wide transcriptional regulation in fast muscle of zebrafish following a single-satiating meal, J. Exp. Biol. 214 (2011) 2125–2139. 10.1242/jeb.053298.

[15] N.I. Bower, I.A. Johnston, Discovery and characterization of nutritionally regulated genes associated with muscle growth in Atlantic salmon, Physiol. Genomics. 42A (2010) 114–130. 10.1152/physiolgenomics.00065.2010.

[16] N.I. Bower, D.G. de la serrana, I.A. Johnston, Characterisation and differential regulation of MAFbx/Atrogin-1 α and β transcripts in skeletal muscle of Atlantic salmon (Salmo salar), Biochem. Biophys. Res. Commun. 396 (2010) 265–271. 10.1016/j.bbrc.2010.04.076.

[17] B.M. Cleveland, J.P. Evenhuis, Molecular characterization of atrogin-1/F-box protein-32 (FBXO32) and F-box protein-25 (FBXO25) in rainbow trout (Oncorhynchus mykiss): Expression across tissues in response to feed deprivation, Comp. Biochem. Physiol. B Biochem. Mol. Biol. 157 (2010) 248–257. 10.1016/j.cbpb.2010.06.010.

[18] M.J.H. Gilbert, T.C. Zerulla, K.B. Tierney, Zebrafish (Danio rerio) as a model for the study of aging and exercise: Physical ability and trainability decrease with age, Exp. Gerontol. 50 (2014) 106–113. 10.1016/j.exger.2013.11.013.

[19] C.-C. Sun, Z.-Q. Zhou, Z.-L. Chen, R.-K. Zhu, D. Yang, X.-Y. Peng, L. Zheng, C.-F. Tang, Identification of Potentially Related Genes and Mechanisms Involved in Skeletal Muscle Atrophy Induced by Excessive Exercise in Zebrafish, Biology. 10 (2021) 761. 10.3390/biology10080761.

[20] W. Xie, M. He, D. Yu, Y. Wu, X. Wang, S. Lv, W. Xiao, Y. Li, Mouse models of sarcopenia: classification and evaluation, J. Cachexia Sarcopenia Muscle. 12 (2021) 538–554. 10.1002/jcsm.12709.

[21] K.-W. Baek, Y.-K. Jung, J.-S. Kim, J.S. Park, Y.-S. Hah, S.-J. Kim, J.-I. Yoo, Rodent Model of Muscular Atrophy for Sarcopenia Study, J. Bone Metab. 27 (2020) 97–110. 10.11005/jbm.2020.27.2.97.

[22] Daya, R. Donaka, D. Karasik, Zebrafish models of sarcopenia, Dis. Model. Mech. 13 (2020) dmm042689. 10.1242/dmm.042689.

[23] C.A. MacRae, R.T. Peterson, Zebrafish as tools for drug discovery, Nat. Rev. Drug Discov. 14 (2015) 721–731. 10.1038/nrd4627.

[24] L. Maves, Recent advances using zebrafish animal models for muscle disease drug discovery, Expert Opin. Drug Discov. 9 (2014) 1033–1045. 10.1517/17460441.2014.927435.

[25] G. Kawahara, J.A. Karpf, J.A. Myers, M.S. Alexander, J.R. Guyon, L.M. Kunkel, Drug screening in a zebrafish model of Duchenne muscular dystrophy, Proc. Natl. Acad. Sci. 108 (2011) 5331–5336. 10.1073/pnas.1102116108.

[26] G.H. Farr, M. Morris, A. Gomez, T. Pham, E. Kilroy, E.U. Parker, S. Said, C. Henry, L. Maves, A novel chemical-combination screen in zebrafish identifies epigenetic small molecule candidates for the treatment of Duchenne muscular dystrophy, Skelet. Muscle. 10 (2020) 29. 10.1186/s13395-020-00251-4.

[27] S.B. Rutkove, S. Callegari, H. Concepcion, T. Mourey, J. Widrick, J.A. Nagy, A.K. Nath, Electrical impedance myography detects age-related skeletal muscle atrophy in adult zebrafish, Sci. Rep. 13 (2023) 7191. 10.1038/s41598-023-34119-6.

