## Supplementary material for "Cold-induced muscle atrophy in zebrafish: Insights from swimming activity and gene expression analysis": Figs. S1-S3

### Slide 1
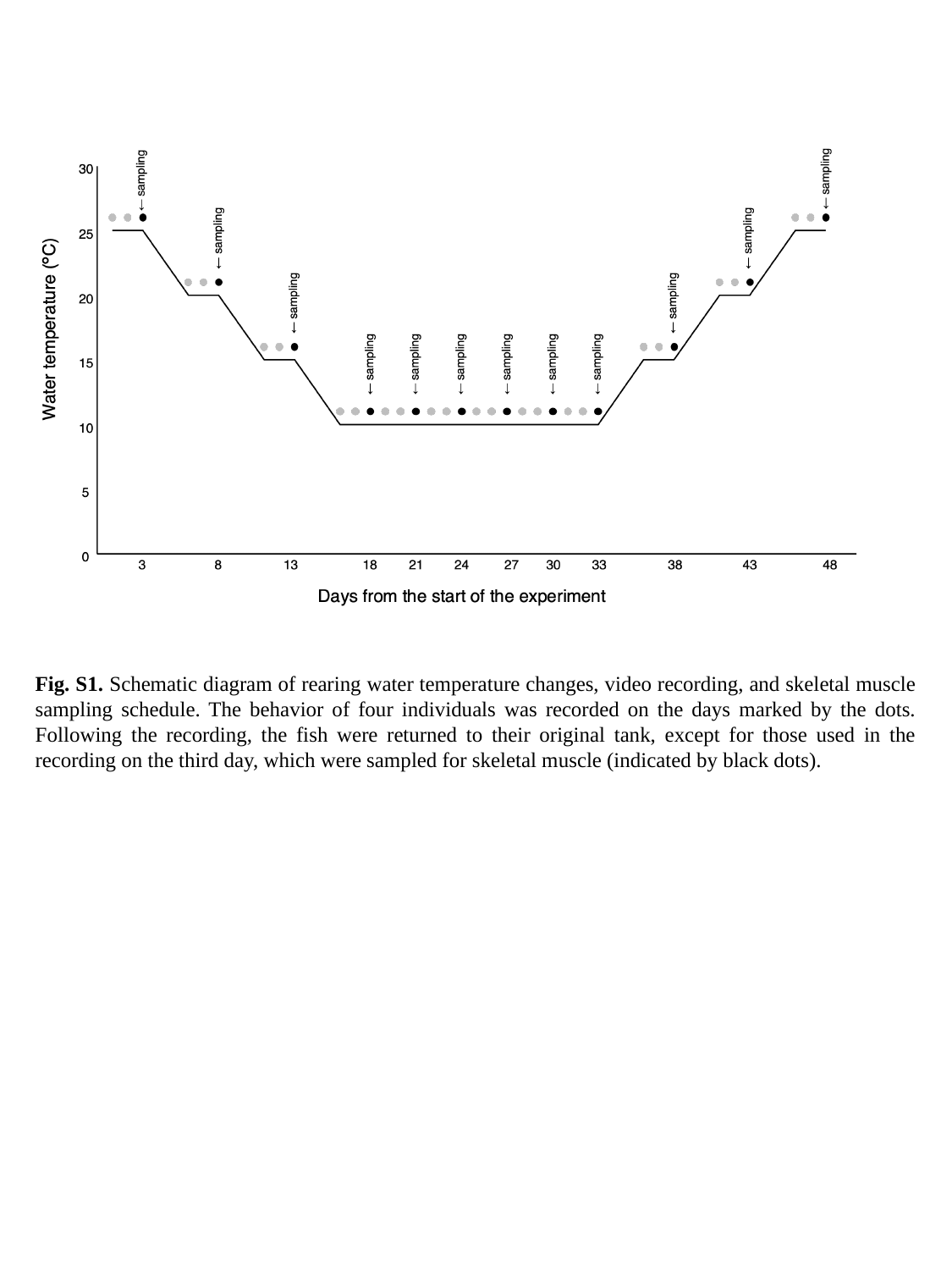

Fig. S1. Schematic diagram of rearing water temperature changes, video recording, and skeletal muscle sampling schedule. The behavior of four individuals was recorded on the days marked by the dots. Following the recording, the fish were returned to their original tank, except for those used in the recording on the third day, which were sampled for skeletal muscle (indicated by black dots).

### Slide 2
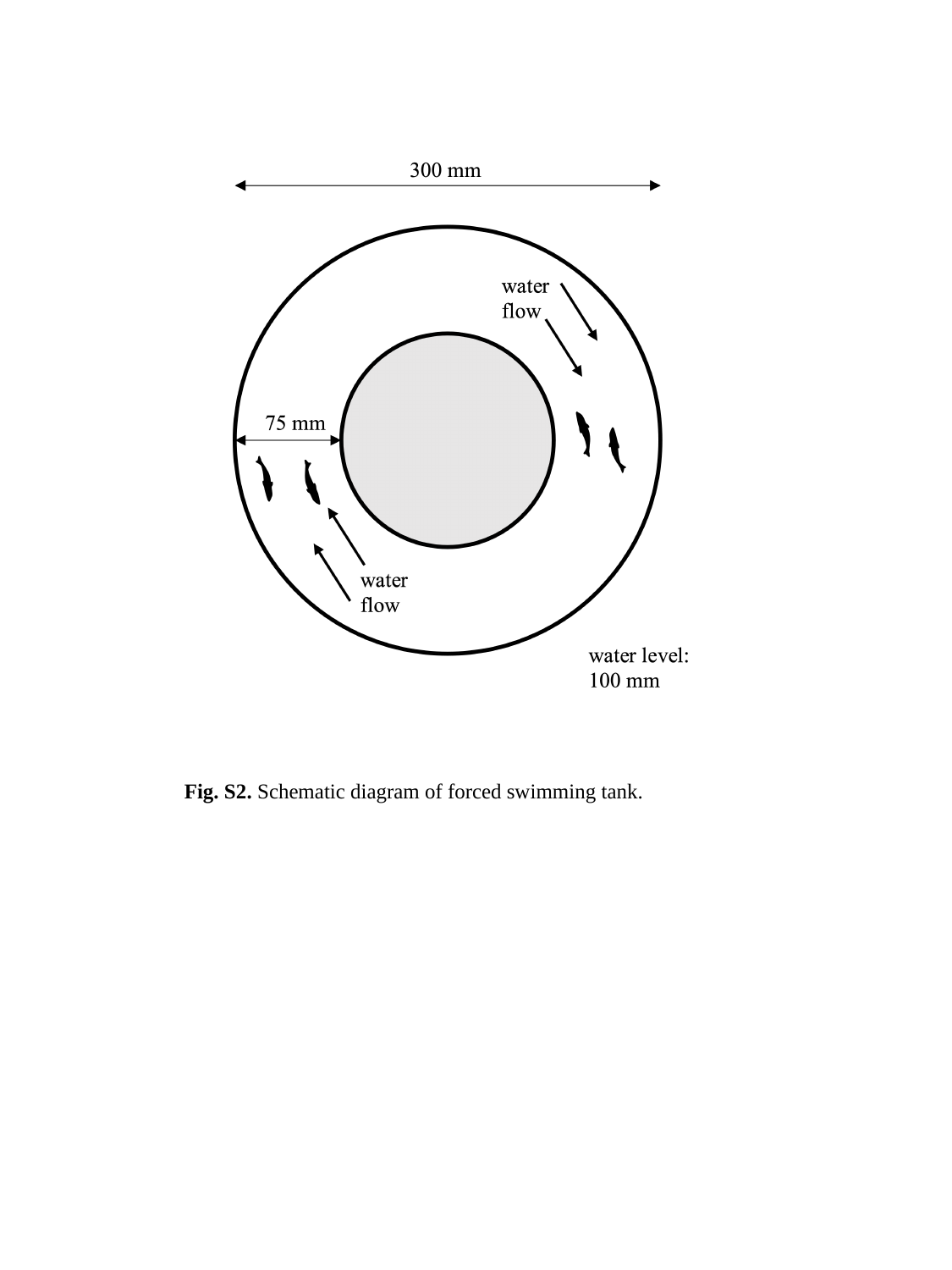

Fig. S2. Schematic diagram of forced swimming tank.

### Slide 3
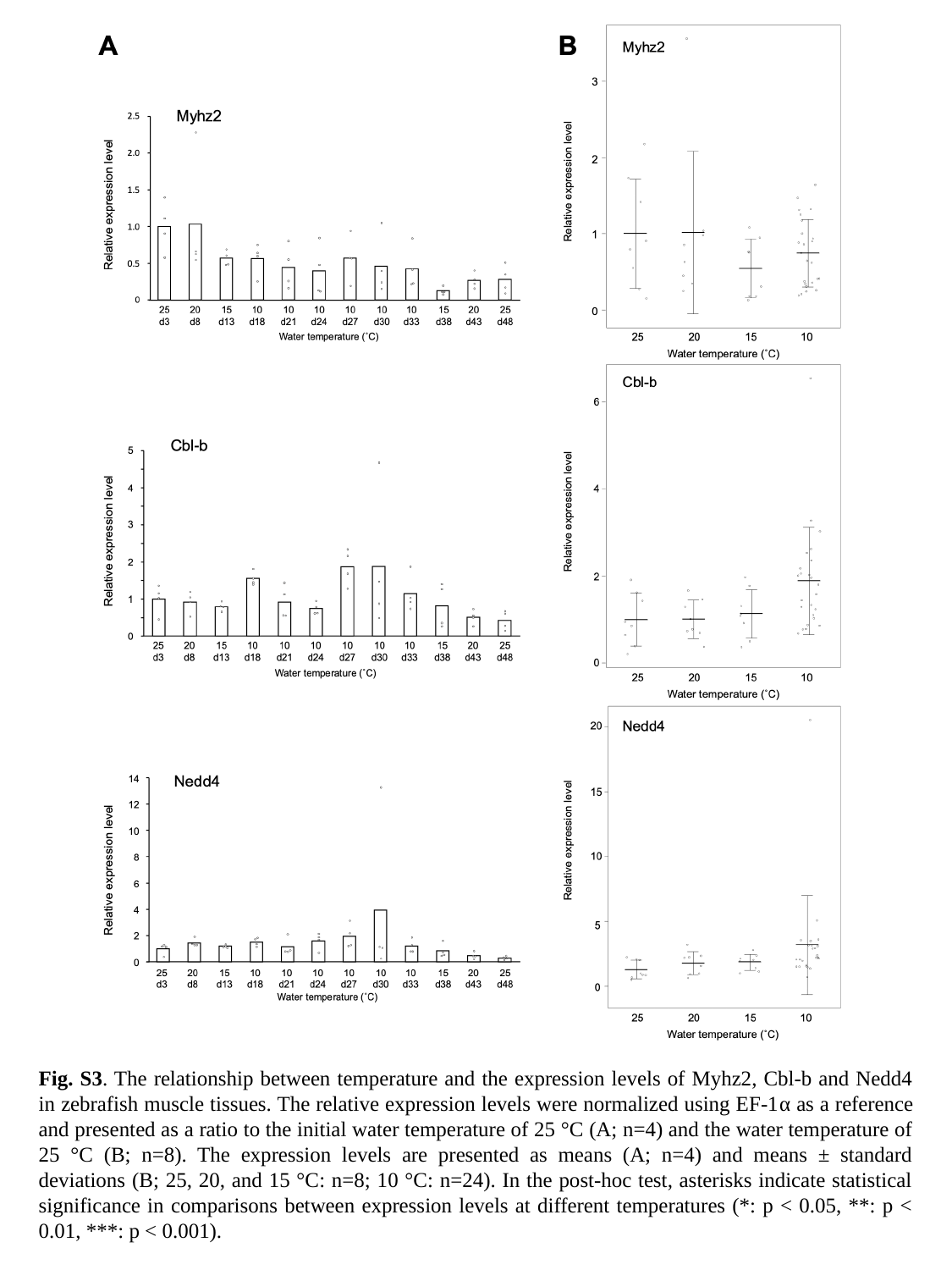

Fig. S3. The relationship between temperature and the expression levels of Myhz2, Cbl-b and Nedd4 in zebrafish muscle tissues. The relative expression levels were normalized using EF-1α as a reference and presented as a ratio to the initial water temperature of 25 °C (A; n=4) and the water temperature of 25 °C (B; n=8). The expression levels are presented as means (A; n=4) and means ± standard deviations (B; 25, 20, and 15 °C: n=8; 10 °C: n=24). In the post-hoc test, asterisks indicate statistical significance in comparisons between expression levels at different temperatures (*: p < 0.05, **: p < 0.01, ***: p < 0.001).
