## Supplementary figures and images for "Cold-induced muscle atrophy in zebrafish: Insights from swimming activity and gene expression analysis"

### Table S1

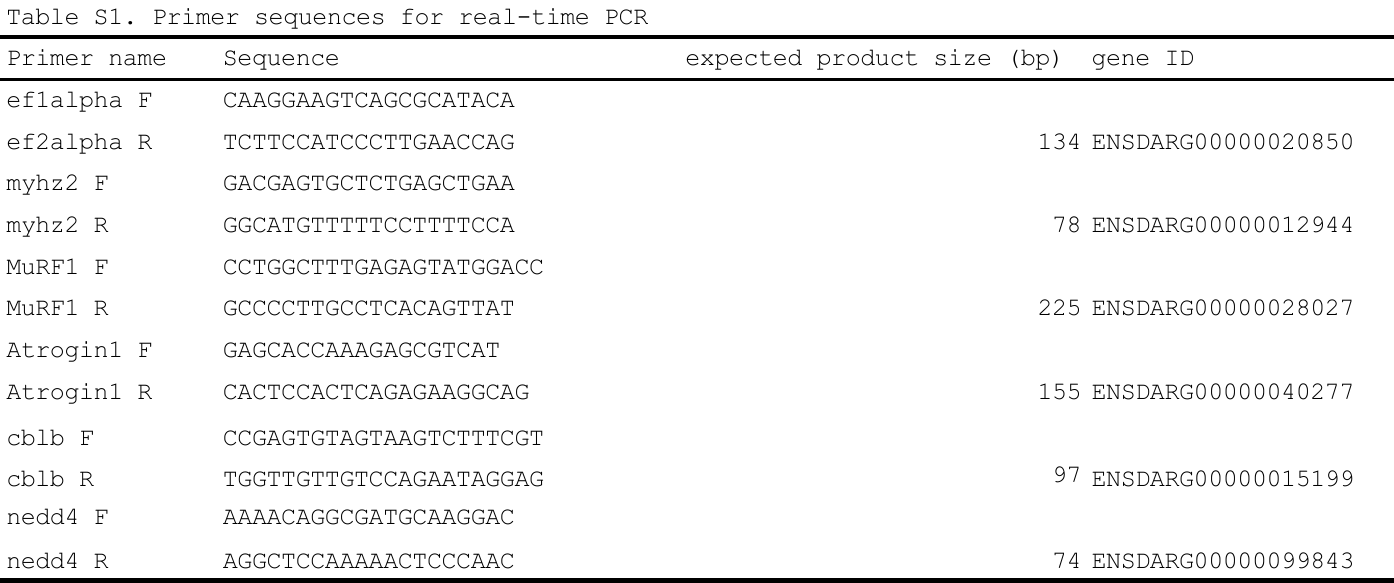
